# Inclusion of dimethyl sulfoxide- dissolved protease inhibitor cocktail in the lysis buffer cuts down the number of proteins resolved by two dimensional gel electrophoresis

**DOI:** 10.1101/2020.05.22.111062

**Authors:** P.P Mahesh, Sathish Mundayoor

**Author notes:** Inter University Centre for Biomedical Research, Kottayam 686009, Kerala, India (Sathish Mundayoor). **E mail:** (Correspondence).

## Abstract

Two dimensional gel electrophoresis (2DE) resolves a mixture of proteins based on both isoelectric point and molecular weight of the individual proteins. Even when we followed a standard protocol for 2DE we got lesser number of proteins focused especially in the basic region of the IPG strip. Since the common troubleshooting measures did not solve the problem we replaced the protease inhibitor cocktail in the lysis buffer with PMSF which resulted in an ideal protein map following 2DE. We also found that the presence of the cocktail results in skewing of the mass spectra of a purified protein which eventually resulted in incorrect identification of the protein by MASCOT search. Later we found that dimethyl sulfoxide, the solvent of protease inhibitor cocktail also resulted in focusing of lesser number of proteins. Addition of 1% dimethyl sulfoxide to bovine serum albumin resulted in lesser sequence coverage for the protein by LC-MS/MS. Also, dimethyl sulfoxide was found to decrease the intensity value of the dominant peptide in a fraction when MALDI-TOF-TOF was done. Even though we do not provide a reason behind our observations, we guess dimethyl sulfoxide might have changed the charge distribution of peptides or proteins resulting in the peculiar observations we had.

## Introduction

Two dimensional gel electrophoresis (2DE) helps to resolve individual proteins from a mixture of proteins based on both the isoelectric point (pI) and molecular weight of proteins. We followed a standard protocol for 2DE of the lysate of THP1 cell line, but we observed that a very few numbers of proteins were resolved and the number of proteins focused in the basic region (pI ˃7 in a pI: 3-10 IPG strip) were less than those focused in the acidic region. This condition made it impossible to compare protein maps of different samples. Therefore we moved on to the conventional troubleshooting measures that are well described in the literature. The troubleshooting included increasing the total protein load, changing the isoelectric focusing conditions, changing the ampholyte concentration, changing the cell lysis conditions and the composition of lysis buffer but none of the measures could solve the observed problem.

## Results and Discussion

When the protease inhibitor cocktail (P8340, Sigma) included in the lysis buffer was withdrawn and replaced with 1mM PMSF we got protein maps with the number of proteins increasing multiple folds (Fig. 1).

**Figure 1.**
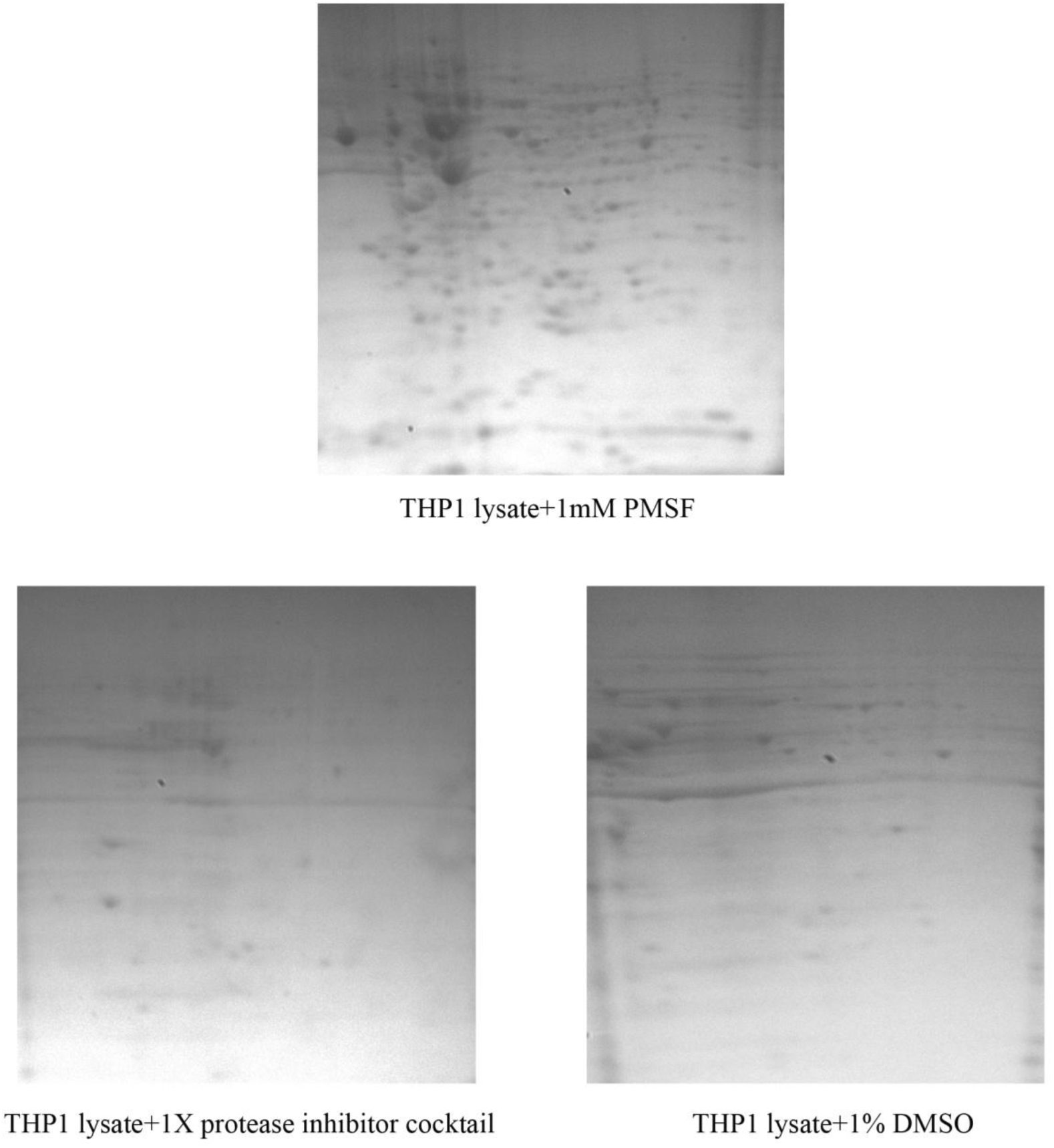
Protease inhibitor cocktail dissolved in DMSO causes missing of proteins from 2DE gels. THP1 lysate containing 1mM PMSF, 1X Protease inhibitor cocktail in DMSO or 1% DMSO was resolved by 2DE

We could also observe that the protein spots picked from 2DE gels made from the lysate containing protease inhibitor cocktail affected the identification of proteins by MALDI-TOF-TOF. Even though the quantity of the proteins picked were higher the corresponding mass spectra were non-ideal with low MOWSE score and slight changes in the MASCOT search parameters resulted in entirely different hits. To confirm this effect we analyzed the mass spectra of purified annexin A2. The purified protein was run on 2DE following treatment with 1X protease inhibitor cocktail, 1mM PMSF or without any treatment. The mass spectrum for the protein in the case of protease inhibitor cocktail treatment was non-ideal and gave entirely different protein hits while Annexin A2 was identified in other cases (Fig. 2).

**Figure 2.**
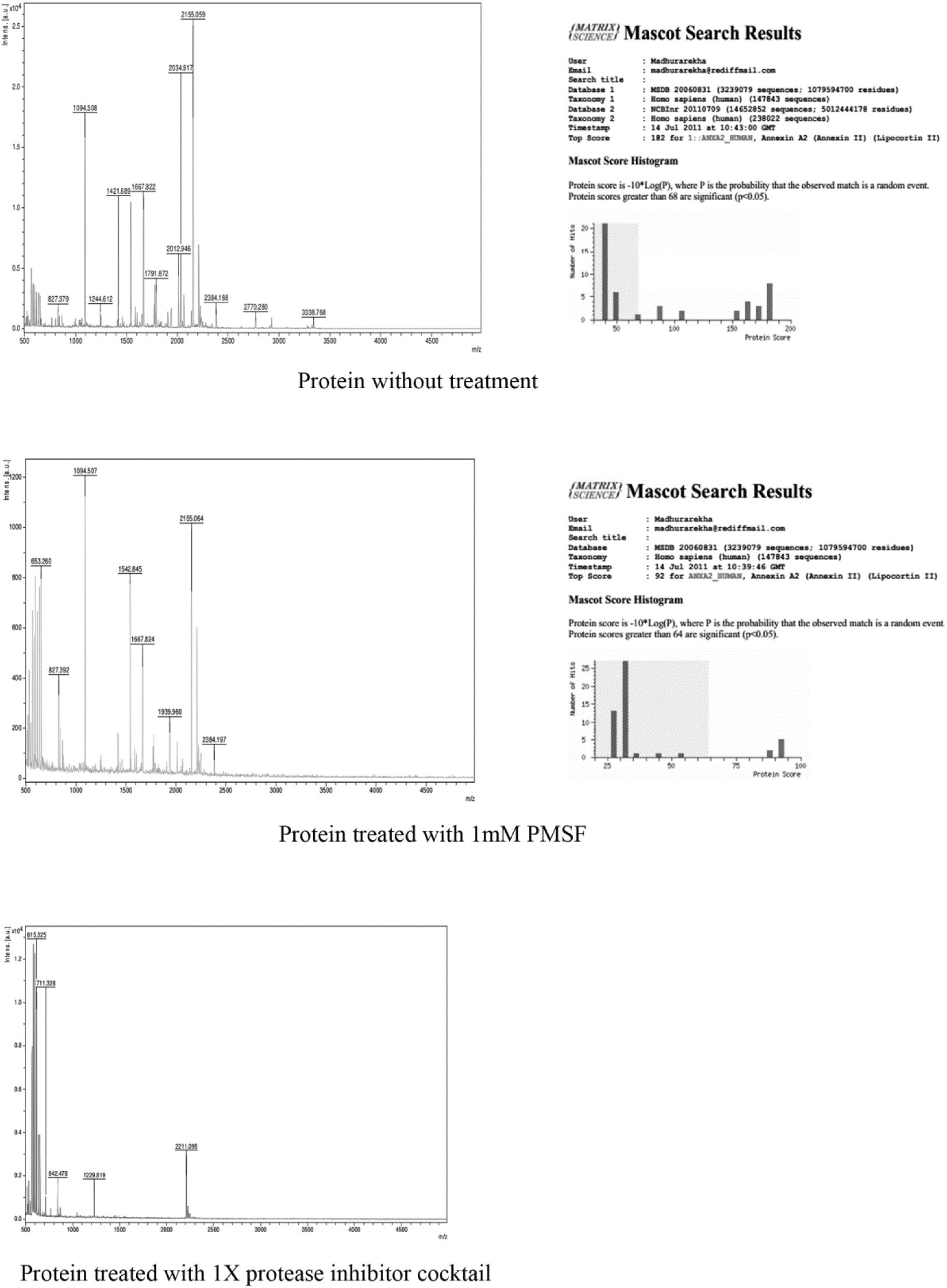
Protease inhibitor cocktail dissolved in DMSO skews the mass spectra of proteins. Purified Annexin A2 treated with 1mM PMSF, Protease inhibitor cocktail dissolved in DMSO or without treatment was run on 2DE and analyzed by MALDI-TOF-TOF.

A study shows that AEBSF, one of the components in the protease inhibitor cocktail, modifies proteins by making a covalent bonding with the amino acid serine and causes a shift in pI of the proteins^1^. This modification causes focusing of the respective proteins in more basic areas of IPG strip. But this finding does not explain our observation of the missing of proteins from 2DE gels. Therefore we assumed that the solvent used in the protease inhibitor cocktail, dimethyl sulfoxide (DMSO), might have an effect. We prepared THP1 lysate and added 1% DMSO (approximately equal to the DMSO concentration when 1X concentration of the protease inhibitor cocktail was used in the lysate), and 2DE was done. 2DE of THP1 lysates containing 1X protease inhibitor cocktail or 1mM PMSF (dissolved in ethanol) were also done for comparison. The 2DE gel of the lysate containing 1% DMSO was more similar to the gel obtained in the case of lysate containing 1X protease inhibitor cocktail with reduced number of proteins compared to the gel obtained in the case of lysate containing PMSF (Fig. 1).

Further, we did 2DE of bovine serum albumin (BSA) treated with 1% DMSO, 1X protease inhibitor cocktail or PMSF and analyzed the mass spectrum by LC-MS/MS followed by tryptic digestion. We could not observe any extra mass corresponding to the molar mass of DMSO on any of the BSA peptides. At the same time it was found that the peptide coverage for BSA in the case of both DMSO and protease inhibitor cocktail were considerably less than in the case of PMSF (Table 1).

**Table 1.** DMSO brings down the sequence coverage for BSA

| Treatment of BSA | Protein score | Average mass | Matched peptides | Digested peptides | Sequence coverage (%) |
| --- | --- | --- | --- | --- | --- |
| 1mM PMSF | 12690.04 | 71289.43 | 49 | 59 | 78.91 |
| 1X protease inhibitor cocktail | 4999.686 | 71289.43 | 37 | 59 | 48.93 |
| 1% DMSO | 402.6649 | 71289.43 | 13 | 59 | 23.23 |

We also did MALDI-TOF-TOF of a synthesized peptide of MW~2540 treated with or without 1% DMSO. The mass spectrum of the peptide fraction (containing a few fragments with different MWs) in water shows the peptide with m/z~2540 having the highest intensity. But the mass spectrum of DMSO-treated peptide shows a peptide with m/z~2578 having the highest intensity while the intensity of m/z~2540 is minimal (Fig. 3). The difference in mass between the two m/z does not correspond to DMSO or any other reagents used for the synthesis of the peptide.

**Figure 3.**
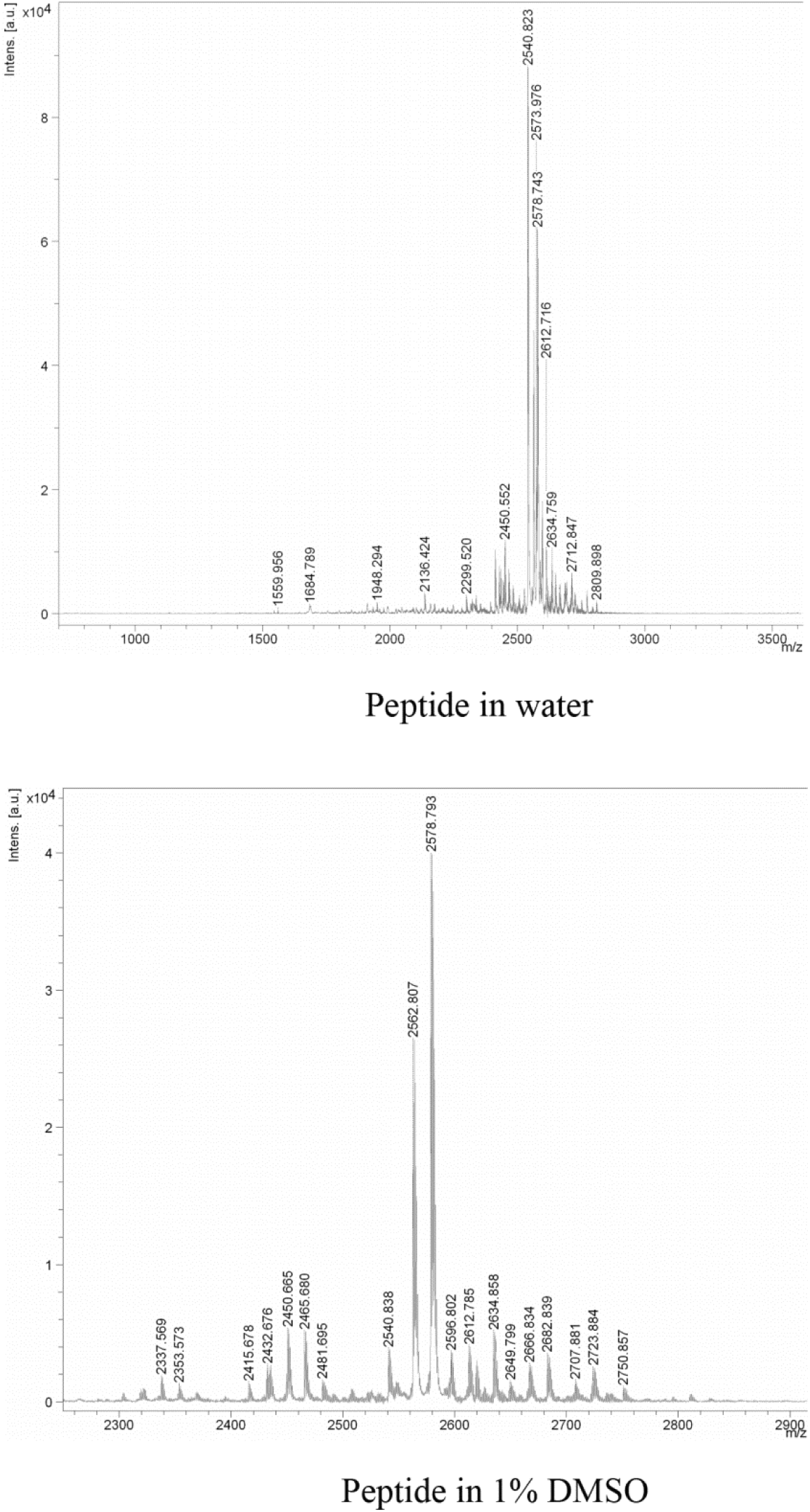
DMSO changes the mass spectrum of the peptide

Earlier studies have reported on the solvent effects of DMSO. A study reports that DMSO destabilizes proteins and shows that low DMSO concentrations influence the ionization process in ESI-MS, resulting in a loss of the protein ion signal intensity as well as a decrease in the basic sites available for protonation on the protein^2^. DMSO was found to increase the basicity of aliphatic amines^3^. When DMSO was added in the peptide mixture, multiple protonation (super charging) of peptides occurs which increases the efficiency of chromatographic separation and ionization^4^. We understand that DMSO changes charge state of peptides and proteins. A change in charge state of proteins may change the pI of proteins and it may make the proteins not to focus within the pI range of an IPG strip during 2DE. Since DMSO changes the mass spectra of proteins it may lead to incorrect identification of proteins by mass spectrometry especially in the case of protein samples with low concentration.

## Materials and methods

### 2D gel electrophoresis and mass spectrometry

Total protein of THP1 cell line was isolated using 2D lysis buffer composed of 8M urea, 1M thiourea and 4% CHAPS along with 1mM PMSF. After 30 minutes of incubation at room temperature the cell lysate was centrifuged at 14000g for 20 min at 18°C and then 300µg of total protein was loaded on a 7cm IPG strip of 3-10 pI range (163-2000, BIO RAD). After isoelectric focusing, the strips were equilibrated and PAGE (15% gel) was run to resolve the proteins. MALDI-TOF-TOF and LC-MS/MS were done in RGCB Proteomics facility and in Vimta Labs, Hyderabad.

## Acknowledgements

This work was supported by intramural fund from RGCB. PPM was supported by a fellowship from Council for Scientific and Industrial Research, Govt of India. Authors thank Dr Ajay Kumar for a purified protein, Dr K Santhosh Kumar for a synthesized peptide and Dr Abdul Jaleel for providing Proteomics facilities.

## Author Contribution statement

PPM and SM designed the study. PPM performed the research. PPM and SM wrote the manuscript together.

